# Expression analysis of defense-related genes in cucumber (*Cucumis sativus* L.) against *Phytophthora melonis*

**DOI:** 10.1101/2020.05.02.073601

**Authors:** Lida Hashemi, Ahmad Reza Golparvar, Mehdi Nasr Esfahani, Maryam Golabadi

**Affiliations:** Department of Agronomy and Plant Breeding, College of Agriculture, Isfahan (Khorasan) Branch, Islamic Azad University, Isfahan, Iran; Department of Agronomy and Plant Breeding, College of Agriculture, Isfahan (Khorasgan) Branch, Islamic Azad University, Isfahan, Iran; Isfahan Center for Agricultural and Natural Resources Research and Education (AREEO), Isfahan, Iran

**Keywords:** *CsWRKY20*, *CsLecRK6*, Damping-off, Genotypes, Gene expression, *LOX1*

## Abstract

*Phytophthora melonis* is the causal agent of damping-off or crown rot, one of the most destructive cucumber diseases that causes severe economic losses in Iran and some other parts of the world. Despite intense research efforts made in the past years, no permanent cure currently exists for this disease. With the aim to understand the molecular mechanisms of defense against *P. melonis*, root collars and leaves of four cucumber genotypes consisting of resistant Ramezz; moderately resistant Baby and very susceptible Mini 6-23 and Extrem, were monitored for quantitative gene expression analysis of five antifungal and/or anti-oomycete genes (*CsWRKY20, CsLecRK6.1, PR3, PR1-1a* and *LOX1*) at three points after inoculation with *P. melonis*. The gene expression analysis indicated that *P. melonis* strongly enhanced the expression of these genes after inoculation in both leaves and root collars. Further, not only the transcript levels of these genes were significantly higher in the resistant and moderately resistance genotypes, but also the time point of the highest relative expression ratio for the five genes was different in the four cucumber genotypes. *CsWRKY20* and *PR3* showed the maximum expression in Ramezz at 48 hours post inoculation (hpi) while *CsLecRK6.1*, and *LOX1* showed the highest expression at 72 hpi. In addition, *PR1-1a* showed the maximum expression in the Baby at 72 hpi. Root collars responded faster than leaves and some responses were more strongly up-regulated in root collars than in leaves. The genes found to be involved in disease resistance in two different organs of cucumber after pathogen infection. The results suggest that increased expression of these genes led to activation of defense pathways and could be responsible for a reduced *P. melonis* colonization capacity in Ramezz and Baby. Overall, this work represents a valuable resource for future functional genomics studies to unravel the molecular mechanisms of *C. sativus*- *P. melonis* interaction.

## Introduction

Cucumber (*Cucumis sativus* L.), a member of the family, Cucurbitaceae, is the fourth most important vegetable crop worldwide (Ren et al. 2009; Sebastian et al. 2010). However, cucumber like other crops is suffering from various pathogens such as different species of Oomycetes. Damping-off disease caused by the hemibiotrophic oomycete, *Phytophthora melonis*, is one of the most severe diseases of the cucurbitaceae, significantly reducing crop yield worldwide (Erwin and Ribeiro 1996; Wu et al. 2014). The main symptoms of cucumbers infected with *P. melonis* are root and root collar rot, stem lesions, foliar blight and fruit rot and finally plant death (McGrath 2001; Hatami et al. 2013; Nasr Esfahani et al. 2012). Damping-off can have a severe economic impact on cucumber from seedling up to fruiting stages. Although the use of disease-resistant genotypes is a key to environmentally friendly and economically sustainable disease control in modern crop production, the employment of genetic resistance to minimize yield losses induced by *P. melonis* remains largely unexplored in cucumber. Up to now, no resistant cultivar has been developed and few reports are available with regard to *Phytophthora* damping-off (Mansoori and Banihashemi, 1982; Nazavari et al., 2016; Hashemi et al., 2019). Therefore, identifying the sources of resistance and studying the genetics underlying resistance to *P. melonis* is pertinent to support cucumber breeding programs.

Plants respond to pathogen attack with a multicomponent defence response that includes synthesis of defense related enzymes like PAL, POX, SOD and CAT, induction a variety of defence genes and enhancement of the cell wall (Van Verk et al., 2009; Anil et al., 2014; Andersen et al., 2018). As most of these defense responses can be monitored at the transcriptional level, gene expression analysis can provide insights into the type of defense mechanism involved in the damping-off disease reaction and cucumber plant pathosystem.

Pathogenesis-related proteins (PRs) are one of the most commonly induced proteins during plant defense mechanism, which have an important role in plant immunity (Van Loon et al. 2006). PR1, PR3 and lipoxygenases (LOX) are strongly induced when plants respond to infection by different types of pathogens. *PR1* gene expression in plant cells is a useful molecular marker for SA-dependent SAR signaling and it could be related to its putative direct antimicrobial action. For example, a basic PR1 protein has been shown to exhibit strong inhibitory activity against the *Phytophthora infestans* in potato (Niderman et al., 1995). whereas PR-3 (endochitinases) is often used as markers for JA-dependent SAR signaling. It hydrolyzes the chitin components of the cell walls of many fungi, such as *R. solani*. Furthermore, the degradation products of the fungal cell wall, especially the oligomers, could serve as resistance elicitors (Rawat et al., 2017). Additionally, lipoxygenase (LOX) pathways are crucial for lipid peroxidation processes during defense responses to infection and limits pathogen growth (Yang et al., 2012). The role of LOX in plant defense against pathogen infection seems to be related to the synthesis of various compounds with signaling functions. In cucumber, the gene expression analysis indicated that *F. oxysporum* strongly enhanced the expression of *PR3, LOX1* and *NPR1*, after inoculation (Pu et al., 2014). Interestingly, some WRKY transcription factors (TFs) were shown to be involved in plant protection responses. Therefore, these genes probably play important roles in combating exogenous pathogens and will provide a basis for further studies of the cloning and functional verification of them and finally, will help us to better understand of the regulatory mechanism of plant resistance to pathogens (Pu et al. 2014; Wang and Bouwmeester 2017). WRKY TFs have been implicated in the regulation of transcriptional reprogramming associated with plant immune responses and genetic evidence demonstrating their significance as positive and negative regulators of disease resistance has accumulated (Noman et al., 2018). For example, in pepper, typical W-boxes were enriched in promoter regions of CaWRKY6 and CaWRKY40, representing involvement of these WRKY TFs in a WRKY network during the regulation of pepper reactions to *Ralstonia solanacearum* inoculation (Ifnan Khan et al., 2018). WRKY TFs positively regulate plant immunity to pathogens associated with H2O2 production, hypersensitive response (HR) mimic cell death and activation of phytohormones mediated signaling pathways. Some of these factors appear to affect the balance between signaling branches promoting SA-dependent and suppressing JA-dependent responses and are required for both basal defense and resistance genes (R-gene) mediated disease resistance against the oomycete (Knoth et al. 2007, Xu et al. 2015). Hussain et al. (2018) showed the role of CaWRKY22 as a positive regulator of pepper immunity against *R. Solanacearum.* WRKY62 positively regulates wheat HTSP resistance to *Puccinia striiformis* f. sp. by differential regulation of SA, JA, ET and ROS-mediated signaling (Wang et al. 2017). In *Paeonia lactiflora*, the PlWRKY65 enhances the resistance to *Alternaria tenuissima* (Wang et al., 2020). Hence, Pathogen infection has been shown to induce strong and rapid induction of WRKY genes from a number of plants. *CsWRKY20* from cucumber was induced by *P. melonis*, may be involved in disease resistance against damping-off (Xu et al. 2015).

The *LecRK* gene families are another group of genes that plays important roles under biotic and abiotic stresses in plants (Wang and Bouwmeester 2017; Boutrot et al. 2017). Recent findings, however, revealed the importance of *LecR*Ks in plant innate immunity (Wang and Bouwmeester 2017; Zhao et al. 2018). In cucumber, CsLecRK6.1 was especially induced by *P. melonis* and *P. capsici* in resistant cultivar (JSH) (Wu et al. 2014; Tingquan et al., 2014). However, the WRKY TFs and LecRKs involved in biotic stresses and disease resistance in cucumber are still largely unknown. However, knowledge about the local (root collars) molecular defense responses compared with systemic (leaves) defenses before and after *P. melonis* inoculation in cucumber is scarce. So far as we know, this is the first study that provides information regarding root collar and leaf gene expression level of *CsWRKY20, CsLecRK6.1, PR3, PR1-1a* and *LOX1* genes in four contrasting genotypes of cucumber upon inoculation with *P. melonis*.

## Materials and methods

### Plant material and Pathogenesis assay

We have recently screened thirty-eight cucumber genotypes, including domestic and exotic hybrids, and inbreeding lines from different seed companies, for damping off resistance to *P. melonis* at two growth stages: seedling (45-dayold seedlings) and maturing (50% flowering) stages (Hashemi et al., 2019). Based on the results of the screening test, Ramezz with high disease resistance, Baby with moderately resistance, Mini 6-23 and Extrem with high susceptibility to *P. melonis* were used throughout the study (Fig 1). The study was carried out in the laboratory of the Agricultural and Natural Resource Research and Education Centre, Isfahan, Iran, in 2018. Cucumber plants of each genotype were cultivated in a single seedling nursery tray filled with substrate of oven-sterilized mixture of sand-peat moss in equal parts in a growth condition at 26– 28 °C with 80% relative humidity. 45-day old seedlings were used for experiments and were inoculated with *P. melonis* isolate, as described by Nasr Esfahani et al. (2012, 2014). The pathogen *P. melonis* was isolated from naturally infected cucumber plants exhibiting post-emergence damping-off and root rot symptoms and identified as *P. melonis* (MH924841). Koch’s postulates were conducted by inoculating onto test cucumber genotypes. Cucumber seedlings were inoculated by drenching with 5 ml of sporangia suspension (106 sporangia/ml) and incubated for two days under saturated moist condition in the greenhouse and then inoculated and non-inoculated treatments grown at 26 ± 2°C, 16 h photoperiod and 65% relative humidity. Cucumber seedlings inoculated with the same amount of sterile distilled water were regarded as the non-inoculated control. Following the inoculation, root collars and leaves of Ramezz, Baby, Mini 6-23 and Extrem were separately collected at four different periods: 0 hour before inoculation, 24, 48 and 72 hpi.

**Fig 1:**
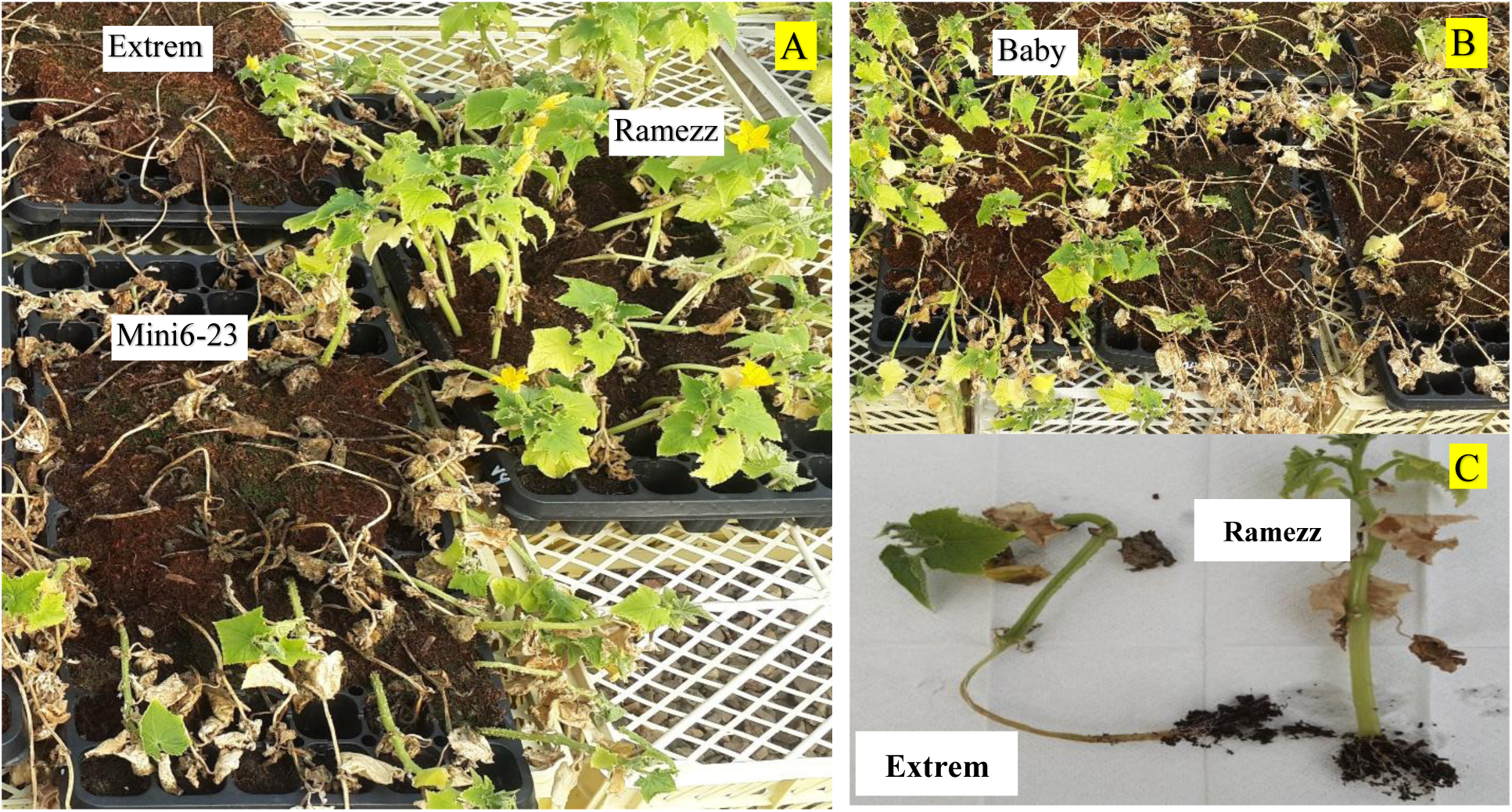
Comparison of damping-off symptom severity between four cucumber genotypes to *P. melonis.* (A) Ramezz (resistant), Extrem and Mini 6-23 (very susceptible), (B) Baby (Moderately resistant), (C) *P. melonis* induced changes in cucumber plants. Images were taken at 7 days’ post inoculation. Severe symptoms (root and collar rot) in very susceptible Extrem was compared with resistant Ramezz. After 4 weeks, very susceptible Extrem inoculated with *P. melonis* showed root and root collar rot symptoms.

### RNA isolation and cDNA synthesis

The infected leaves and root collars from three biological replicates (one replicate representing a pool of three plants) were flash-frozen after collection and separately ground with a pestle and mortar. Total RNA was extracted using AccuZolTM kit (Bioneer, Korea), according to the manufacturer’s instructions. The genomic DNA in the total RNA was eliminated by digestion with RNase-free DNase (Thermo Scientific) following the manufacturer’s protocol. The quantity and purity of total RNA was evaluated using a Nanodrop NP80 spectrophotometer (Implen Germany). The cDNA was synthesized from 5 μg of total RNA using the cDNA Synthesis commercial kit (YTA, Yekta Tajhiz Azma, Iran) according to the manufacturer’s instructions.

### Primer design and relative quantification of gene expression

Gene-specific primers were designed using Oligo7 software (Rychlik, 2007); on *Cucumis sativus* L. gene sequences at the National Center for Biotechnology Information (NCBI; www.ncbi.nlm.nih.gov). The primer sequences and their amplicon characteristics for each gene are presented in Table1. The specificity of primers designed was confirmed by performing the PCR reaction on synthesized cDNA and genomic DNA (To confirm the accuracy of the PCR).

### qRT-PCR

The mRNA expression levels of 5 genes were analyzed by quantitative real-time PCR (qRT-PCR) using SYBR Green PCR Master Mix (2X) (Biofact, South Korea) according to the manufacturer’s protocols. All analysis was carried out on StepOne™ 48-well Real-Time PCR System (Applied Biosystems). The qRT-PCR was performed with preliminary hold at 95 °C for 15 min, 45 cycles of 95 °C for 15 s, annealing temperature depending on the primer Tm for 25 s and 72 °C for 40 s (Table 1). The experiment was performed with three biological replicates for each sample and two technical replicates. Primer efficiencies (E) for all primers and Ct values were determined using LinRegPCR software version 2012.3 (Ruijter et al., 2009) and the Pfaffl method was used to calculate the relative gene expression (Pfaffl, 2001). The cucumber *actin* gene (XM_004147305.2) was used as the internal reference for the normalization of expression level (Wan et al. 2010; Castellano et al. 2016).

### Growth measurements

After 7 days of *P. melonis* inoculations, plants were harvested to measure some growth parameters. Root fresh weights, root dry weight, shoot fresh weight and shoot dry weight were expressed in grams per plant. Root volume was measured using the method of changes in the water volume. The root and Collar diameter (in mm) was measured by using a digital caliper with accuracy of 0.01 mm. In addition, root length was expressed in cm.

### Statistical analysis

The qRT-PCR results were analyzed using a completely randomized factorial design with three replications and data analysis was performed using the StepOneTM Software v2.3. Analysis of variance (ANOVA) was with SAS (ver. 9.1, SAS Institute, Cary, NC) and means comparison was with Least Significant Difference (LSD) (p<0.05) (Li et al. 2018).

## Result

### Effect of damping-off on the growth parameters of cucumber plants inoculated with *P. melonis*

*Phytophthora melonis* inoculation decreased fresh and dry weights of roots and shoots, root volume and collar diameter between cucumber genotypes significantly. This reduction was greater in high susceptible Mini6-23 and Extrem (Table 2). No difference in these plant growth parameters was observed between the inoculated Ramezz, compared to the healthy control (Table 2). A strong significant negative correlation between the percentage of disease incidence caused by *P. melonis* and the fresh weights of roots (r^2^ = −0.65), dry weights of roots (r^2^ = −0.83), root volume (r^2^ = – 0.68), and collar diameter (r^2^ = −0.70) were determined. There was also a moderate negative correlation between the percentage of disease incidence and root diameter (r2 = −0.57).

### Expression Patterns of *CsWRKY20* and *CsLecRK6.1* genes in leaves after infection with *P. melonis*

As shown in Fig1, the transcripts of the genes *CsLecRK6.1* and *CsWRKY20* responding to different time points were detected in resistant, moderately resistant and high susceptible genotypes but at different levels. In the Ramezz and Baby, *CsLecRK6.1* transcript levels was strongly up-regulated as early as 24 hpi (3.02 and 2.32 fold) which gradually increased and reached a peak at 72 hpi (7.86 and 7.94 fold), respectively (Fig2.a). In contrast, high susceptible Mini 6-23 and Extrem showed consistent and low expression of *CsLecRK6.1* until 72 h after infection. The gene, *CsWRKY20*, had increased expression levels between 0 to 48 hpi and then declined from 48 hpi to 72 hpi, although in Ramezz and Baby was still higher than the transcript expression in Mini 6-23 and Extrem (Fig2.b). *CsWRKY20* transcript accumulation in leaves of Ramezz showed the strongest, 6.78-fold enhancement at 48 hpi compared with the control (Fig2.b). The differential expressions of these candidate transcripts may lead to the difference in resistance level among different genotypes and tissues.

**Fig 2:**
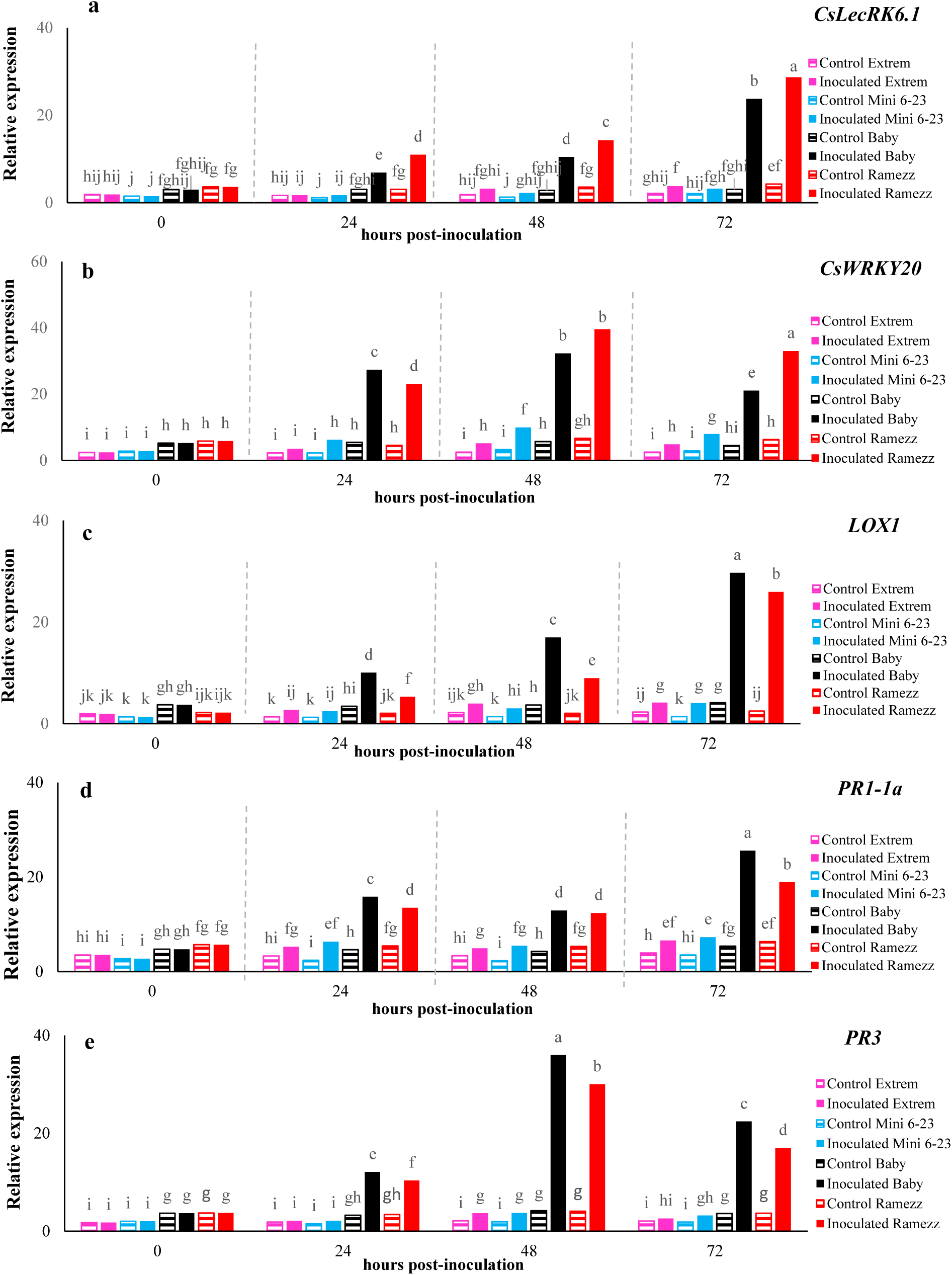
Relative transcript levels of *CsLecRK6.1, CsWRKY20, PR1-1a, PR-3* and *LOX1* genes in leaves of non-inoculated (dark horizontal) and inoculated (solid fill) resistant cucumber genotype: Ramezz (red), moderately resistant genotype: Baby (balck) and very susceptible genotypes: Mini 6-23 (blue) and Extrem (pink) at 0 hours before and 24, 48 and 72 hours post inoculation with *P. melonis*. Data represent the means ± SD (n = 3) of three biological replicates. Different letters denote significant differences at P <0.05.

### Expression Patterns of *WRKY20* and *CsLecRK6.1* genes in root collars after infection with *P. melonis*

Similar to that observed in leaves, transcript levels of *CsLecRK6.1* were significantly elevated in root collars of Ramezz and Baby at 24 hpi and thereafter (Fig3.a). While, *CsLecRK6.1* transcript levels were significantly lower in the root collars of Mini 6-23 and Extrem relative to Ramezz and Baby. So, the resistant Ramezz showed the strongest, over nine-fold enhancement of *CsLecRK6.1* transcription at 72 hpi compared with the control (Fig3.a). There was significant positive correlation between relative *CsLecRK6.1* expression level in leaves and root collars (r = 0.85**). The gene, *CsWRKY20*, had increased expression levels until 48 hpi in Ramezz and Baby, decreasing thereafter at 72 hpi. In contrast, high susceptible genotypes showed no significant difference in gene expression (Fig3.b). Also, significant positive correlations were recorded for relative *CsWRKY20* expression level in leaves and root collars (r = 0.98**).

**Fig 3:**
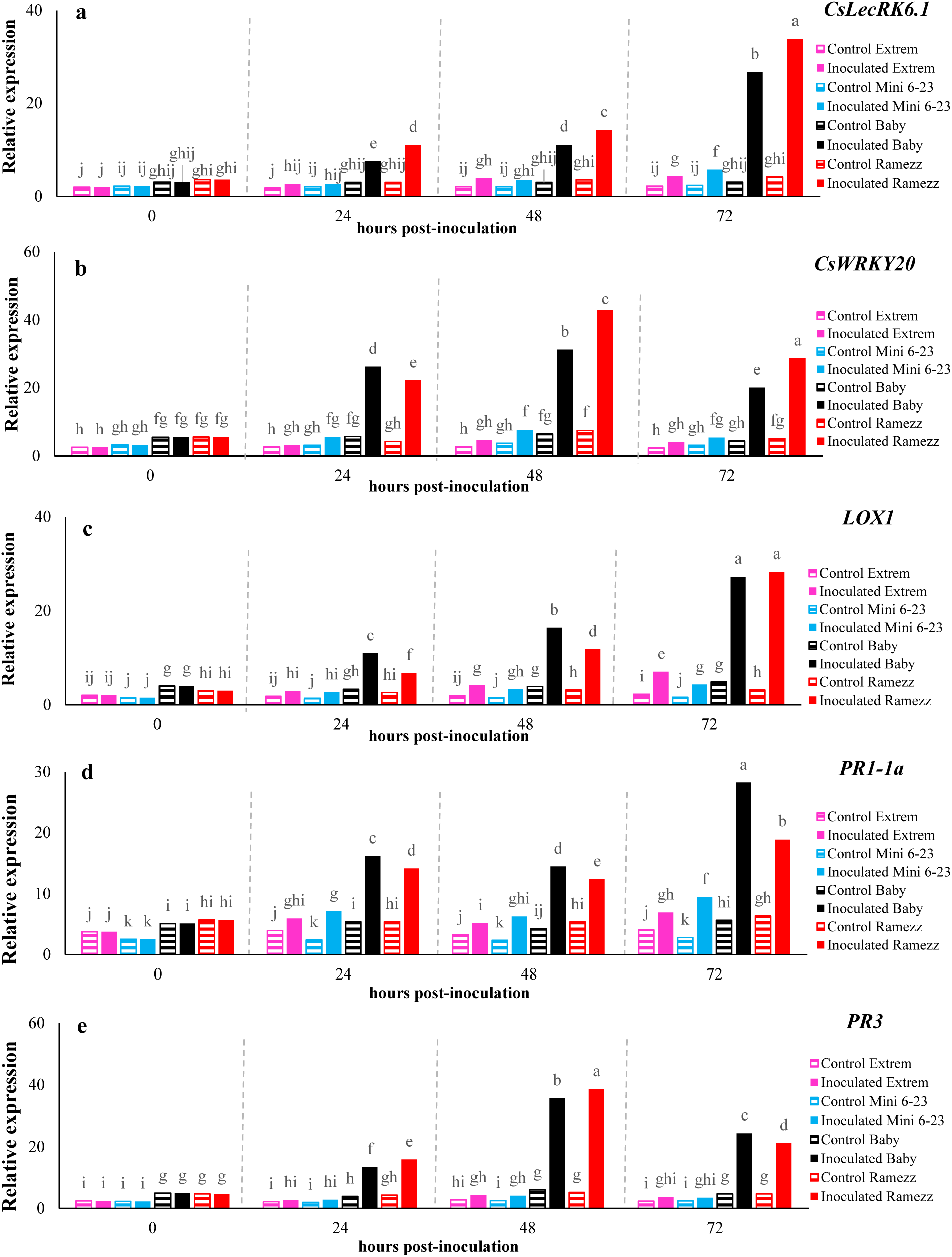
Relative transcript levels of *CsLecRK6.1, CsWRKY20, PR1-1a, PR-3* and *LOX1* genes in root collars of non-inoculated (dark horizontal) and inoculated (solid fill) resistant cucumber genotype: Ramezz (red), moderately resistant genotype: Baby (balck) and very susceptible genotypes: Mini 6-23 (blue) and Extrem (pink) at 0 hours before and 24, 48 and 72 hours post inoculation with *P. melonis*. Data represent the means ± SD (n = 3) of three biological replicates. Different letters denote significant differences at P <0.05.

### Expression Patterns of *PR3, PR1-1a* and *LOX 1* genes in Leaves after infection with *P. melonis*

The *PR3* was up-regulated in *P. melonis*-infected leaves of all genotypes in the first 48 hpi followed by a decrease at 72 hpi, while the overall expression remained lower in the high susceptible genotypes (Mini 6-23 and Extrem) when compared to resistant Ramezz and moderately resistant (Ramezz and Baby). The expression level of *PR3* in the leaves of Ramezz increased as much as 8.01 fold at 48 hpi compared with the control, while in Baby increased 9.72 fold which decreased in both Ramezz and Baby at 72 hpi (Fig2.e). The transcript levels of *PR1-1a* were increased in leaves of cucumber plants at 24 hpi and thereafter (Fig2. d). The expression levels of *PR1-1a* in leaves of Ramezz at 24, 48 and 72 hpi up-regulated 2.37, 2.18 and 3.32 fold, respectively, in Baby by 3.34, 2.73 and 5.39 fold, respectively, compared to the control. Interestingly, the gene *PR1-1a* expression was significantly higher in Baby than Ramezz at all the time points. The transcript levels of the gene *LOX1* increased (24, 48 and 72 hpi) in leaves of the resistant and moderately resistance genotypes, higher in Ramezz than Baby (Fig2.c). Although the relative expression of *LOX1* in Mini 6-23 and Extrem was significant at 48 and 72 hpi, but the expression level was lower than Ramezz and Baby. The *LOX1* expression in Ramezz and Baby was up-regulated about 11.81 and 8.36 fold in the leaves, respectively, compared to the control (Fig2.c). The higher level of transcript for *LOX, PR1-1a* and *PR3* in Ramezz and Baby suggests that they are likely responsible for a large part of resistance in cucumber.

### Expression Patterns of *PR3, PR1-1a* and *LOX1* genes in root collars after infection with *P. melonis*

The *PR3* was up-regulated in *P. melonis*-infected leaves of all genotypes in the first 48 hpi followed by a decrease at 72 hpi (Fig3.e). The highest expression level of *PR3* existed in root collars of Ramezz rather than other genotypes, at 48 hours after inoculation, while in the leaves, it occurred for Baby. The expression of *PR1-1a* gene was significantly increased in *P. melonis*-infected root collars of all genotypes at 24, 48 and 72 hpi, while the overall expression remained lower in Mini 6-23 and Extrem when compared to Baby and Ramezz at all the time points. *PR1-1a* transcript was gradually up-regulated thereby reaching a peak at 72 hpi (Fig3.d). Moreover, the expression level of *PR1-1a* in the two tissues (root collars and leaves) was basically the same. The gene *LOX1* expression was strongly and simultaneously up-regulated in leaves and root collars with a maximum at 72 hpi (Fig3.c). The expression levels of *LOX1* was increased markedly from 24 to 72 hpi and reached a maximum that was 11.82 fold higher than the control (Fig3.c). There were significant positive correlations between the expression levels of these genes in root collars and leaves.

## Discussion

With the aim to explore biological processes involved in the mechanism of resistance/susceptibility of *C. sativus* against the Oomycete, *P. melonis*, we chose to undertake an analysis of expression levels of five candidate defense-related genes in leaves and root collars of selected cucumber genotypes that possibly contribute to the establishment of damping-off resistance in the cucumber plant. Therefore, four cucumber genotypes, which according to the recent screening test results showed resistance (Ramezz), moderately resistance (Baby) and high susceptibility (Mini 6-23 and Extrem) to damping-off, were used to analyze the transcriptional activity of *CsWRKY20, CsLecRK6.1, PR3, PR-1a* and *LOX1* genes during infection produced by *P. melonis.* Patterns of gene expression can provide important clues to the analysis of involvement of defense-related genes in defense processes because it is believed that genes that share the same expression pattern are likely to be connected with regard to their functions. Moreover, comparing the expression patterns of these genes in root collars and leaves of four cucumber genotypes with different degrees of resistance in response to *P. melonis*, helps in the analysis of the differences and commonalities between these two tissues in response to damping-off. Moreover, the resistant Ramezz and moderately resistant Baby showed much earlier and higher expression pattern compared to the high susceptible genotypes, Mini 6-23 and Extrem, under the pathogen stress. The higher activities of defense enzymes such as PAL and POX, and higher level of H2O2 in infected resistant Ramezz and moderately resistant Baby tissue compared to the high susceptible genotype Mini 6-23 and Extrem may explain theirs high level of resistance. The contrasting degrees of susceptibility of these cucumber genotypes to damping-off represents a good opportunity to investigate their specific molecular responses in order to better understand underlying mechanisms governing the resistance or susceptibility to *P. melonis.* This study revealed obvious similarity between leaves and root collars in their response to damping-off. Moreover, the transcripts of these five genes accumulated significantly at a greater level in Ramezz and Baby than in Mini 6-23 and Extrem upon inoculation with *P. melonis* and probably being involved in early responses against *P. melonis*.

Increasing evidence suggests critical role of WRKY TFs in regulating defense related genes and Producing an increased resistance against infection (Finatto et al. 2018). In addition, some WRKY TFs act as positive regulators of SA-dependent defense responses and positively regulate *PR* expression (Jiang et al., 2014; Liu et al. 2018). In the current study, the expression of *CsWRKY20* gene from *Cucumis sativus* was induced after infection by *P. melonis* and a significant accumulation of *CsWRKY20* transcripts was observed in Ramezz and Baby in the present study. So the plants constitutively expressing *CsWRKY20* were more resistant to damping-off. Our finding is in accordance with previous studies which have demonstrated the association of WRKY TFs in resistance response against Oomycete phytopathogens. Xu et al. (2015) predicted that *CsWRKY20* may be involved in Oomycete disease resistance of JSH (resistant cucumber cultivar) against *P. melonis* by JA and (or) SA signaling pathway(s). The *WRKY40* positively regulated the expression of defense-related genes of chickpea and enhanced disease resistance against *Pseudomonas syringae* (Chakraborty et al. 2018). Another study has shown, the *SpWRKY3* may enhance tomato resistance to tomato late blight (*Phytophthora infestans*) as a positive regulator (Cui et al. 2018). Therefore, this is not surprising considering the fact that *CsWRKY20* in cucumber might has a similar role during interaction between cucumber and *P. melonis.* Further research through the development and analysis of loss of function mutants for *CsWRKY20* gene is needed to clarify its role in cucumber signaling in response to *P. melonis* infection.

Previous study showed that *CsLecRK6.1* may play an important role in disease resistant against *P. melonis* in JSH resistant cucumber (Wu et al. 2014). Additionally, several studies highlight the significant and outstanding role of *LecRKs* in plant innate immunity on several aspects that make them fascinating as potential resistance components in plant (Wang and Bouwmeester 2017; Zhao et al. 2018). Accordingly, we selected *CsLecRK6.1* as candidate and study results demonstrate that *CsLecRK6.1* responded to *P. melonis* in resistant and moderately resistant genotypes and a significant accumulation of gene transcripts was observed in the Ramezz and Baby for all the time point after inoculation. Bouwmeester et al. (2011) demonstrated that expression of *LecRK-I.9* was induced by inoculating with non-host and avirulent pathogens in Arabidopsis and subsequently enhanced resistance to *Phytophthora brassicae* by overexpression of *LecRK-I.9*. A recent study by Wang et al. (2018) showed that *LecRK-V* confers broad-spectrum resistance to powdery mildew in wheat and SA pathways contribute to the enhanced resistance to the virulent powdery mildew fungus. However, the expression information of *CsLecRK6.1* gene will be useful for further investigating the function of this gene under various biotic stress conditions.

We also focused on the expression pattern of some PR genes that are reliable defence markers in cucumber, such as PR1-1a, a marker of the SA pathway, PR3 encoding chitinases. PR proteins are accumulated in various parts of normal tissues and are induced by pathogen infection and improve the defensive capacity of plants. Members of the PR-1 and PR-3 families of proteins have direct activities against Oomycete pathogens (Sudisha et al. 2012). Sterol-auxotroph pathogens such as the Phytophthora are particularly sensitive to PR-1 families and plants with enhanced PR-1 expression are particularly well protected against Oomycete (Gamir et al., 2017; van den Berg et al. 2018). The transcripts of *PR1-1a* was significantly up-regulated in both leaves and root collars of inoculated cucumber plants. Additionally, the gene *PR1-1a* expression was significantly higher in moderately resistant and resistant genotype than high susceptible genotypes.

It is thought that Chitinase1, which is encoded by *PR3* and can hydrolyze chitin, contributes to the defense of plants (Rawat et al. 2017). Study of cucumber and nonpathogenic *F. oxysporum* CS-20 interaction have shown a rapid response to pathogen inoculation at the initial stage, 24 hpi, and that it increased again at 48 hpi and then reduced again at 72 hpi (Pu et al. 2014). In this study, we find similar patterns of *PR-3* expression. Indeed, *PR-3* was considered as a key gene involved in activation of the defence response and *PR-3* (endochitinases) can hydrolyze the pathogen cell walls. So, *PR-3* was up-regulated in all genotype in the first 48hrs.

The significant and active accumulation of *PR1-1a* and *PR-3* in Ramezz and Baby suggests a crucial role of these transcripts in direct defense mechanism of cucumber against the *P. melonis*. Therefore, it may be hypothesized that the difference in susceptibly between the four genotypes could be explained, at least partly, by the differential expression of genes involved in anti-oomycete defense.

In response to *P. melonis*, Ramezz and baby show greater levels of induction of lipoxygenase gene, *LOX1* coding for enzyme of the biosynthesis of JA. Previous studies support the role of LOX genes in plant defense against fungal and oomycete infection (Hwang and Hwang, 2010; Maschietto et al. 2015). Pathogen-induced *LOX* transcript accumulation has been reported in a number of plants, e.g. in cucumber after inoculation with *F. oxysporum* CS-20 (Pu et al. 2014), in tobacco after infection with *P. parasitica* var. nicotianae (Rancé et al. 1998) and in potato infected by *P. infestans* (Göbel et al. 2002). Although, this interaction is not the same, we found a similar profile in our experiment, in which *LOX1* showed activation after inoculation by *P. melonis*. Consequently, the up-regulation of *PR1-1a* and *PR-3* on one side and the up-regulation of *LOX1* on the other side, suggests a much intense activation of SA and JA pathways in Ramezz and Baby than in Mini 6-23 and Extrem, which probably promote the tolerance to *P. melonis*. Pu et al. (2014) showed, maximum expression level of *PR3, NPR1* and *LOX 1* genes were observed at 24, 48 and 72 hours after infection by *Fusarium oxysporum*. So Based on the results of Pu et al. (2014), we selected 24/48/72hpi as sampling times and our result confirmed the results of previous research on cucumber against *F. oxysporum*. Additionally, our results indicated that *CsWRKY20* and *CsLecRK6.1* as well as *PR* genes (*PR3, PR-1a*) responded quickly in leaves and root collars to *P. melonis* stress, suggesting that these responses might be central components of the induced immune system through JA and SA signaling pathway. Surprisingly, among the 5 defence-related genes monitored in infected leaves and root collars, the *LOX1* and *CsLecRK6.1* gene were highly up-regulated in the inoculated cucumber plants, respectively. In addition, the similar response of these genes expression may indicate that same control mechanisms of these genes may exist in leaves and root collars. Additionally, infection of cucumber with *P. melonis* resulted in a significant decrease in fresh and dry weights of roots and aerial parts, root volume and collar diameter, although the magnitude of these decrease depended upon the genotypes. Previous research showed that all Phytophthora species were able to cause damage to the root and shoot growth parameters (Aghighia et al., 2016; Ruiz Gómez et al., 2018).

Our results showed that the defense response due to *P. melonis* inoculation varied in resistance, moderately resistance and high susceptible genotypes of cucumber and the difference between resistance and susceptibility depends on genetic variation in resistance. Based on the fact that the expression of *PR3, PR1-1a, LOX1, CsLecRK6.1* and *CsWRKY20* genes in infected plants was significantly different from the control at 0 hour before inoculation with *P. melonis*, it is suggested that these genes mostly participate in defensive response. Furthermore, these genes showed different levels of transcript accumulation in investigated genotypes and the higher levels of expression of these genes was observed in Ramezz and Baby. All these results strongly indicated that under *P. melonis* stress conditions leaves and root collars of different cucumber genotypes employed same mechanisms to cope with damping-off.

## Conclusion

Although, this is a preliminary study on *C. sativus*- *P. melonis* interaction, but could be useful to understand defense mechanism of cucumber. Additionally, the study of these gene expression profiles will further promote the functional research of cucumber plant defense responses and signaling pathways in the future. However, more research is needed to detect the differences and gain an insight into the more detailed characteristics of these genes and interactions of *P. melonis* effectors with the genes under study.

## Acknowledgement

Thanks go to Plant Protection Research Division, Isfahan Center for Agricultural and Natural Resources Research and Education (AREEO), Isfahan, Iran and also, Plant Protection Research Institute, Tehran, Iran, for providing facilities to run the project.

## References

Aghighia, S., Burgessa, T. I., Scottbc, J. K., Calveraand, M., Hardy, G. E. St. J. (2016). Isolation and pathogenicity of Phytophthora species from declining Rubus anglocandicans. Plant Pathology. 65(3), 451–461.

Alves, M. S., Dadalto, S. P., Goncalves, A.B., DeSouza, G. B., Barros, V. A., & Fietto, L. G. (2014). Transcription factors functional protein-protein interactions in plant defense responses. Proteomes, 2, 85–106.

Andersen, E. J., Ali, S., Byamukama, E., Yen, Y., Nepal, M. P. (2018). Disease Resistance Mechanisms in Plants. Genes, 9(7): 339. https://doi.org/10.3390/genes9070339.

Anil, K., Das, S. N, Podile, A. R. (2014). Induced defense in plants: a short overview. The Proceedings of the National Academy of Sciences, India, Section B: Biological Sciences, 84, 669–679.

Boutrot, F., & Zipfel, C. (2017). Function, discovery, and exploitation of plant pattern recognition receptors for broad-spectrum disease resistance. Annual Review of Phytopathology, 55, 257–286.

Bouwmeester, K., de Sain, M., Weide, R., Gouget, A., Klamer, S., Canut, H., et al. (2011). The lectin receptor kinase LecRK-I.9 is a novel *Phytophthora* resistance component and a potential host target for a RXLR effector. PLoS Pathogens, 7, e1001327. Pmid: 21483488.

Castellano, M., Pallas, V., & Gomez, G. (2016). A pathogenic long noncoding RNA redesigns the epigenetic landscape of the infected cells by subverting host Histone Deacetylase 6 activity. New Phytologist, 211(4), 1311–22.

Chakraborty, J., Ghosh, P., Sen, S., & Das, S. (2018). Epigenetic and transcriptional control of chickpea WRKY40 promoter activity under *Fusarium* stress and its heterologous expression in Arabidopsis leads to enhanced resistance against bacterial pathogen. Plant Science, 276, 250–267.

Cui, J., Xu, P., Meng, J., Li, J., Jiang, N., & Luan, Y. (2018). Transcriptome signatures of tomato leaf induced by *Phytophthora infestans* and functional identification of transcription factor SpWRKY3. Theoretical and Applied Genetics, 131(4), 787–800.

Erwin, D. C., & Ribeiro, O. K. (1996). Phytophthora Disease Worldwide. APS Press. St. Paul, MN, pp: 562.

Finatto, T., Viana, V. E., Woyann, L. G., Busanello, C., Maia, L., Oliveira, A. C. (2018). Can WRKY transcription factors help plants to overcome environmental challenges? Genetics and molecular biology, 41(3), 533–544. https://doi.org/10.1590/1678-4685-GMB-2017-0232.

Gamir, J., Darwiche, R., Van’t Hof, P., Choudhary, V., Stumpe, M., Schneiter, R., Mauch, F. (2017). The sterol-binding activity of PATHOGENESIS-RELATED PROTEIN 1 reveals the mode of action of an antimicrobial protein. The Plant Journal, 89(3), 502–509.

Göbel, C., Feussner, I., Hamberg, M., Rosahl, S. (2002). Oxylipin profiling in pathogeninfected potato leaves. Biochimica et Biophysica Acta, 1584, 55–64.

Hashemi, L., Golparvar, A. R., Nasr Esfahani, M., Golabadi, M. (2019). Correlation between cucumber genotype and resistance to damping-off disease caused by Phytophthoramelonis, Biotechnology & Biotechnological Equipment, 33:1, 1494–1504, DOI: 10.1080/13102818.2019.1675535

Hatami, N., Aminaee, M. M., Zohdi, H., & Tanideh, T. (2013). Damping-off disease in greenhouse cucumber in Iran, archives of phytopathology and plant protection, 46(7), 796–802.

Hussain, A., Li, X., Weng, Y., Liu, Z., Ashraf, M. F., Noman, A., Yang, S., Ifnan, M., Qiu, S., Yang, Y., Guan, D., He, S. (2018). CaWRKY22 Acts as a Positive Regulator in Pepper Response to *Ralstonia Solanacearum* by Constituting Networks with CaWRKY6, CaWRKY27, CaWRKY40, and CaWRKY58. International journal of molecular sciences, 19(5), 1426. https://doi.org/10.3390/ijms19051426.

Hwang, I. S., & Hwang, B. K. (2010). The pepper 9-lipoxygenase gene CaLOX1 functions in defense and cell death responses to microbial pathogens. Plant physiology, 152(2), 948–967. https://doi.org/10.1104/pp.109.147827.

Ifnan Khan, M., Zhang, Y., Liu, Z., Hu, J., Liu, C., Yang, S., Hussain, A., Furqan Ashraf, M., Noman, A., Shen, L., Yang, F., Guan, D., He, S. (2018). CaWRKY40b in Pepper Acts as a Negative Regulator in Response to *Ralstonia solanacearum* by Directly Modulating Defense Genes Including CaWRKY40. International Journal of Molecular Sciences, 19(5):1403. doi: 10.3390/ijms19051403. PMID: 29738468; PMCID: PMC5983674.

Jiang, Y., Duan, Y., Yin, J., Ye, S., Zhu, J., Zhang, F., Lu, W., Fan, D., Luo, K. (2014). Genome-wide identification and characterization of the Populus WRKY transcription factor family and analysis of their expression in response to biotic and abiotic stresses. Journal of Experimental Botany, 65:6629–6644.

Knoth, C., Ringler, J., Dangl, J.L., Eulgem, T. (2007) Arabidopsis WRKY70 is required for full RPP4-mediated disease resistance and basal defense against *Hyaloperonospora parasitica*. Molecular Plant-Microbe Interactions, 20, 120–128.

Li C, Ren Y, Jiang S, et al (2018). Effects of dietary supplementation of four strains of lactic acid bacteria on growth, immune-related response and genes expression of the juvenile sea cucumber *Apostichopus japonicus* Selenka. Fish and shellfish immunology 74; 69–75.

Liu, Q., Li, X., Yan, S., Yu, T., Yang, J., Dong, J., Zhang, S., Zhao, J., Yang, T., Mao, X., Zhu, X., & Liu, B. (2018). OsWRKY67 positively regulates blast and bacteria blight resistance by direct activation of PR genes in rice. BMC Plant Biology, 18, 257.

Mansoori B, Banihashemi Z. Evaluating cucurbit seedling resistance to *Phytophthora drechsleri*. Plant Dis. 1982; 66(1):373–376.

Maschietto, V., Marocco, A., Malachova, A., & Lanubile, A. (2015). Resistance to *Fusarium verticillioides* and fumonisin accumulation in maize inbred lines involves an earlier and enhanced expression of lipoxygenase (LOX) genes. Journal of Plant Physiology, 188, 9–18.

McGrath, M. T. (2001). Vegetable MD online: *Phytophthora* blight of cucurbits. Cooperative Extension, New York State, Cornell University. Online publication.

Nasr Esfahani, M., Chatraee, M., Shafizadeh, S., & Jalaji, S. (2012). Evaluation of resistance of cucurbit and cucumber cultivars to *Phytophthora drechsleri* in Greenhouse. Iranian Seed and Plant Improvement Journal, 28, 407–417.

Nasr Esfahani M, Nasehi A, Rahmanshirazi P, et al. Susceptibility assessment of bell pepper genotypes to crown and root rot disease. Arch Phytopathol Plant Protect. 2014;47(8):944–953.

Nazavari, K., Jamali, F., Bayat, F., & Modarresi, M. (2016). Evaluation of resistance to seedling damping-off caused by *Phytophthora drechsleri* in cucumber cultivars under greenhouse conditions. Biological Forum, 8, 54–60.

Niderman T, Genetet I, Bruyere T, Gees R, Stintzi A, Legrand M, et al. (1995). Pathogenesis-related PR-1 proteins are antifungal—isolation and characterization of 3 14-kilodalton proteins of tomato and of a basic PR-1 of tobacco with inhibitory activity against *Phytophthora infestans*. Plant Physiology, 108, 17–27.

Noman, A., Liu, Z., Aqeel, M., Zainab, M., Khan, MI., Hussain, A., Ashraf, MF., Li, X., Weng, Y., He, S. (2017). Basic leucine zipper domain transcription factors: the vanguards in plant immunity, Biotechnology Letters, 39 (12), 1779–1791.

Pfaffl, M. W. (2001). A new mathematical model for relative quantification in real-time RT-PCR. Nucleic acids research, 29(9), e45.

Pu, X., Xie, B., Li, P., Mao, Z., Ling, J., Shen, H., Zhang, J., Huang, N., & Lin, B. (2014). Analysis of the defense-related mechanism in cucumber seedlings in relation to root colonization by nonpathogenic *Fusarium oxysporum* CS-20. FEMS Microbiology Letters, 355(2), 142–51.

Rancé, I., Fournier, J., & Esquerré-Tugaye, M. T. (1998) The incompatible inter-action between *Phytophtora parasitica* var *nicotianae* race 0 and tobacco is suppressed in transgenic plants expressing antisense lipoxygenase se-quences. Proceedings of the National Academy of Sciences of the United States 95: 6554–6559.

Rawat, S., Ali, S., Mittra, B., & Grover, A. (2017). Expression analysis of chitinase upon challenge inoculation to Alternaria wounding and defense inducers in *Brassica juncea*. Biotechnology Reports, 13, 72–79.

Ren, Y., Zhang, Z., Liu, J., Staub, J. E., Han, Y., et al. (2009). An Integrated Genetic and Cytogenetic Map of the Cucumber Genome. PLoS ONE, 4(6), e5795. doi: 10.1371/journal.pone.0005795.

Ruijter, J. M., Ramakers, C., Hoogaars, W. M., Karlen, Y., Bakker, O., van den Hoff, M. J., & Moorman, A. F. (2009) Amplification efficiency: linking baseline and bias in the analysis of quantitative PCR data. Nucleic Acids Research, 37(6), e45.

Ruiz Gómez, F.J., Pérez-de-Luque, A., Sánchez-Cuesta, R., Quero, J.L., Navarro Cerrillo, M.N., 2018. Differences in the Response to Acute Drought and *Phytophthora cinnamomi* Rands Infection in Quercus ilex L. Seedlings. Forests. 9, 634.

Rychlik, W. (2007). OLIGO 7 Primer Analysis Software. In: Yuryev A. (eds) PCR Primer Design. Methods in Molecular Biology™, vol 402. Humana Press.

Sebastian, P., Schaefer, H., Telford, I. R., & Renner, S. S. (2010). Cucumber (*Cucumis sativus*) and melon (*C. melo*) have numerous wild relatives in Asia and Australia, and the sister species of melon is from Australia. Proceedings of the National Academy of Sciences of the United States of America, 107, 14269–14273.

Sudisha, J., Sharathchandra, R., Amruthesh, K., Kumar, A., & Shetty, H.S. (2012). Plant Defense: Biological Control, in: Pathogenesis Related Proteins in Plant Defense Response (pp. 379–403). New York: Springer.

Tingquan, W., Rui, W., Xiaomei, X., Xiaoming, H., Baojuan, S., Yujuan, Z., Zhaojuan, L., Shaobo, L., & Yu’e, L. (2014). *Cucumis sativus* L-type lectin receptor kinase (*CsLecRK*) gene family response to *Phytophthora melonis, Phytophthora capsici* and water immersion in disease resistant and susceptible cucumber cultivars. Gene, 549(2), 214–22.

Van Loon, L. C., Rep, M., & Pieterse, C. M. J. (2006). Significance of inducible defense-related proteins in infected plants. Annual Review of Phytopathology, 44, 135–162.

Van Verk, M. C., Gatz, C., Linthorst, H. J. M. (2009). Transcriptional regulation of plant defense responses. Advances in Botanical Research, 51, 397–438.

van den Berg, N., Mahomed, W., Olivier, N. A., Swart, V., & Crampton, B. G. (2018). Transcriptome analysis of an incompatible *Persea americana*-*Phytophthora cinnamomi* interaction reveals the involvement of SA- and JA-pathways in a successful defense response. PLOS ONE, 13(10), e0205705.

Wang, J., Tao, F., Tian, W., Guo, Z., Chen, X., Xu, X., Shang, H., Hu, X. (2017). The wheat WRKY transcription factors TaWRKY49 and TaWRKY62 confer differential high-temperature seedling-plant resistance to *Puccinia striiformis* f. sp. tritici. PloS one, 12(7), e0181963. https://doi.org/10.1371/journal.pone.0181963.

Wang, Y., & Bouwmeester, K. (2017). L-type lectin receptor kinases: New forces in plant immunity. PLoS Pathogens, 13(8), e1006433.

Wang, Z. K., Cheng, J. Y., Fan, A. Q., Zhao, J., Yu, Z. Y., Li, Y. B., Zhang, H., Xiao, J., Muhammad, F., et al. (2018). LecRK-V, an L-type lectin receptor kinase in *Haynaldia villosa*, plays positive role in resistance to wheat powdery mildew. Plant Biotechnology Journal, 16, 50–62.

Wang, X., Li, J., Guo, J., Qiao, Q., Guo, X., Ma, Y. (2020). The WRKY transcription factor PlWRKY65 enhances the resistance of *Paeonia lactiflora* (herbaceous peony) to *Alternaria tenuissima*. Horticulture Research, 7, 57. https://doi.org/10.1038/s41438-020-0267-7.

Wu, T., Wang, R., Xu, X., He, X., Sun, B., Zhong, Y., Liang, Z., Luo, S., & Lin, Y. (2014). *Cucumis sativus* L-type lectin receptor kinase (*CsLecRK*) gene family response to *Phytophthora melonis, Phytophthora capsici* and water immersion in disease resistant and susceptible cucumber cultivars. Gene, 549, 214–222.

Xu, X., Wang, R., Chao, J., Lin, Y., Jin, Q., He, X., Luo, S., & Wu, T. (2015). The expression patterns of *Cucumis sativus* WRKY (*CsWRKY*) family under the condition of inoculation with *Phytophthora melonis* in disease resistant and susceptible cucumber cultivars. Canadian Journal of Plant Science, 95, 1121–1131.

Yang, X. Y., Jiang, W. J., & Yu, H. J. (2012). The expression profiling of the lipoxygenase (LOX) family genes during fruit development, abiotic stress and hormonal treatments in cucumber (Cucumis sativus L.). International journal of molecular sciences, 13(2), 2481–2500. https://doi.org/10.3390/ijms13022481.

Zhao, T., Wang, J., Zhang, B., & Hou, X. (2018). Genome-Wide Analysis of Lectin Receptor-Like Kinases in Tomato (*Solanum lycopersicum*) and Its Association with the Infection of Tomato Yellow Leaf Curl Virus. Plant Molecular Biology Reporter, 36: 429–438.

